# Thermal cycling protects SH-SY5Y cells against hydrogen peroxide and β-amyloid-induced cell injury through stress response mechanisms involving Akt pathway

**DOI:** 10.1101/2019.12.19.877373

**Authors:** Wei-Ting Chen, Yu-Yi Kuo, Guan-Bo Lin, Chueh-Hsuan Lu, Hao-Ping Hsu, Yi-Kun Sun, Chih-Yu Chao

## Abstract

Neurodegenerative diseases (NDDs) are becoming a major threat to public health, according to the World Health Organization (WHO). The most common form of NDDs is Alzheimer’s disease (AD), boasting 60-70% share. Although some debates still exist, excessive aggregation of β-amyloid protein (Aβ) and neurofibrillary tangles has been deemed one of the major causes for the pathogenesis of AD. A growing number of evidences from studies, however, have suggested that reactive oxygen species (ROS) also play a key role in the onset and progression of AD. Although scientists have had some understanding of the pathogenesis of AD, the disease still cannot be cured, with existing treatment only capable of providing a temporary relief at best, partly due to the obstacle of blood-brain barrier (BBB). The study was aimed to ascertain the neuroprotective effect of thermal cycle hyperthermia (TC-HT) against hydrogen peroxide (H_2_O_2_) and Aβ-induced cytotoxicity in SH-SY5Y cells. Treating cells with this physical stimulation beforehand significantly improved the cell viability and decreased the ROS content. The underlying mechanisms may be due to the activation of Akt pathway and the downstream antioxidant and prosurvival proteins. The findings manifest significant potential of TC-HT in neuroprotection, via inhibition of oxidative stress and cell apoptosis. It is believed that coupled with the use of drugs or natural compounds, this methodology can be even more effective in treating NDDs.

## Introduction

According to the World Health Organization (WHO), the number of dementia-induced deaths more than doubled between 2000 and 2016, making it the 5th leading cause for deaths worldwide in 2016, up from 14th place in 2000. Among the various forms of dementia such as Alzheimer’s disease (AD) and Parkinson’s disease (PD), AD is the most common one, accounting for 60–70% of the cases. Typically, AD exhibits such features as deposition of cortical plaques caused by excessive aggregation of β-amyloid protein (Aβ) and neurofibrillary tangles, and progressive brain degeneration and deterioration of cognitive function among elderly people. Although the exact mechanism of AD pathogenesis remains unknown, it is believed that oxidative stress and activation of free radicals, induced by Aβ aggregation, play a key role in AD pathogenesis [1].

Reactive oxygen species (ROS) are reactive chemical species containing oxygen, which are generated as natural byproduct of oxygen metabolism. ROS play important roles in cell signalling and homeostasis, and their concentrations in cells are subtlety regulated by various antioxidant compounds and enzymes. However, with cells under continuing environmental stress (e.g. ultraviolet, inflammatory cytokines, or environmental toxins), the imbalance between prooxidants and antioxidants may cause chronic oxidative stress. Accumulation of ROS may cause cell death, accelerate cell ageing, or induce age-related diseases [2]. More and more research evidences have suggested that ROS plays a central role in the onset and progression of AD [3]. Therefore, the protection of neural cells against oxidative damage may be a potential strategy to treat AD. Several in vitro or in vivo studies have explored the function of antioxidant and antiapoptotic drugs in ameliorating AD [4–6] but the approach is time-consuming and costly, plus the safety concern, which limits the use of these drugs in AD treatment. Moreover, the blood-brain barrier (BBB) dampens the efficacy of these drugs, since over 98% of small molecule drugs and ~100% of large molecule drugs can not pass the BBB [7]. Therefore, a non-drug treatment may be more suited to AD patients.

Scientists have long been interested in the profound effects of heat on cells, and have utilized it in various types of thermotherapeutical applications such as physiotherapy, urology, and cardiology [8]. One promising and effective thermal therapy is the treatment of cancer by hyperthermia (HT) [9]. HT is used to kill cancer cells directly or to potentiate the cytotoxicity of radiation and certain chemotherapy drugs [10]. The ROS level increased by HT treatment has been identified to play an important role as an intracellular mediator of HT-induced cell death [11]. On the contrary, it has also been reported that heat shock (HS) will induce many cellular defense, including the antioxidant effect. For example, Tchouagué *et al* demonstrated that HS-generated ROS is involved in induction of cellular defense molecules Prxs, GSH and G6PD through Nrf2 activation [12]. Mustafi *et al* also showed that heat stress upregulates the HSP70 and MnSOD levels through ROS and p38MAPK [13]. In addition to the thermal treatment, the beneficial effects of light treatment were also reported in literatures. The review article by Hamblin summarized some pre-clinical studies and clinical trials by light therapy for brain disorders [14]. The physical stimulation, therefore, holds great potential for AD or other neurodegenerative diseases (NDDs).

The study employed a special thermal therapy, applied at high and low temperatures alternately to achieve an effect similar to antioxidant and antiapoptotic drugs. In our previous study, we used this novel method, thermal cycle hyperthermia (TC-HT) for the treatment of pancreatic cancer, and found that this physical stimulation greatly improved the anticancer effect of propolis and polyphenols on PANC-1 cells without the heat-induced side effect [15, 16]. Traditional HS or HT employs continuous heating to achieve curative effect, but it is likely to cause neuron damage since damage to the central nervous system occurs within few minutes of exposure to 42°C [17]. Nevertheless, the feature of TC-HT is that it can avoid the damage caused by HT, which is crucial for neuroprotection.

In this study, we applied the TC-HT strategy to human neural cell line SH-SY5Y, which has been extensively used in research on neurodegenerative damage in vitro, and examined the prosurvival effect of TC-HT on preventing oxidative damage induced by hydrogen peroxide (H_2_O_2_) and Aβ. The results showed that subjection of the cells to heat at high and low temperatures alternately beforehand ameliorated the H_2_O_2_ and Aβ-induced cytotoxicity in SH-SY5Y cells significantly. It was found that TC-HT not only performed superior protective effect than the traditional HS but also avoided thermal damage caused by continuous heating. Examination of the underlying mechanism also showed that TC-HT could activate specific neuroprotective proteins. These findings indicate that TC-HT is a promising thermal therapy, which sheds light on novel treatment for AD or other NDDs.

## Materials and methods

### Cell culture and treatment

The human neuroblastoma SH-SY5Y cells purchased from Bioresource Collection and Research Center (Hsinchu, Taiwan) were maintained in MEM/F-12 mixture containing 10% fetal bovine serum (HyClone; GE Healthcare Life Sciences) and 1% penicillin-streptomycin, supplemented with 1mM sodium pyruvate and 0.1 mM non-essential amino acids. The cells were cultured at 37°C in a humidified incubator composed of 5% CO_2_. SH-SY5Y cells were seeded in 96-well plates overnight. Cells were pretreated with HT or TC-HT by the Thermal Cycler (Applied Biosystems; Thermo Fisher Scientific, Inc.) relative to the control at room temperature (RT). After the treatment, the cells were maintained in a 37°C incubator for 4 h. Subsequently, the pretreated cells were induced to undergo apoptosis by adding H_2_O_2_ to the culture medium. Cell viability was measured 24 h after the H_2_O_2_ treatment. As for AD disease model, the cells were treated with 25 or 50 µM Aβ_25-35_ protein solution. The Aβ stock solution was prepared by solubilizing the Aβ_25-35_ peptide (Sigma-Aldrich; Merck KGaA) in sterile deionized water to a concentration of 1 mM and then incubated at 37°C for 3 days to allow self-aggregation before treatment. The TC-HT therapy was applied 4 h before the Aβ treatment for the pretreatment group and 1 h after Aβ treatment for the post-treatment group, and the cell viability was determined 4 days after the treatment. For the Akt inhibitor experiment, 12 or 25 µM LY294002 (Cell Signaling Technology, Inc.) was treated 1 h before the TC-HT.

### Cell Viability Assay

The cell viability was determined by 3-(4,5-dimethylthiazol-2-yl)-2,5-diphenyltetrazolium bromide (MTT) (Sigma-Aldrich; Merck KGaA) assay. In brief, the culture medium was replaced with MTT solution (0.5 mg/mL in MEM/F12) and incubated at 37°C for 4 h. Equal volume of the solubilizing buffer (0.01 M HCl and 10% SDS) was added to dissolve the formazan crystals. The 96-well plates were analyzed on a FLUOstar OPTIMA microplate reader (BMG Labtech, Ltd.) at 570 nm, and background absorbance at 690 nm was subtracted.

### Lactate dehydrogenase (LDH) assay

The assay measures a stable enzyme LDH, which is released into the cell culture medium when cell membranes are damaged. LDH levels were measured using a Cytoscan LDH cytotoxicity assay kit (G-Biosciences; Geno Technology Inc.) according to the manufacturer’s instructions. Briefly, the cells were seeded in 96-well plates and then incubated with H_2_O_2_ with or without thermal cycle pretreatment. For analysis, 50 µL culture mediums from all wells were transferred to a new 96-well plate, and 50 µL of reaction mixtures were added to each well and incubated at 37°C for 20 min. The absorbance was measured at 490 nm using a microplate reader (BMG Labtech, Ltd.).

### ROS level detection

ROS levels of cells were detected using the fluorescent dye dihydroethidium (DHE) (Sigma-Aldrich; Merck KGaA). Cells pretreated with TC-HT or HT at 42.5°C high temperature setting for 8 cycles or 2 h, respectively, were challenged with H_2_O_2_. After the treatment of H_2_O_2_ for 24 h, cells were washed with PBS, and then incubated with 5 µM DHE dye for 20 min at 37°C in the dark. The fluorescence intensity emitted by DHE was measured by flow cytometry in the PE channel, and ROS levels were expressed as mean fluorescence intensity for comparison.

### Mitochondrial membrane potential (MMP) measurement

The MMP was determined by flow cytometry using 3,3'-dihexyloxacarbocyanine iodide (DiOC_6_(3)) (Enzo Life Sciences International Inc.). DiOC_6_(3) is a lipophilic cationic fluorescent dye which allows estimation of the percentage of cells with low MMP. Cells were pretreated with TC-HT or HT 4 h before the H_2_O_2_ treatment. After the treatment of H_2_O_2_ for 24 h, cells were harvested and suspended at a density of 1 × 10^6^ cells/mL in 1 µM DiOC_6_(3) dye working solution. After incubation at 37°C for 15 min, DiOC_6_(3) intensity was analyzed by flow cytometry in the FITC channel.

### Western blot analysis

The protein expression levels of SH-SY5Y cells were investigated by western blot analysis. Cells treated with TC-HT, HT, or H_2_O_2_ were harvested and lysed in RIPA lysis butter (EMD Millipore). The cells were harvested within 24 h after H_2_O_2_ treatment, and the timing for different protein collections was adjusted based on previous literatures [18–20]. For HSP70 and HSP105, cells were lysed 16 h after H_2_O_2_ treatment. For p-Akt, Akt, Nrf2, CREB, IDE, PSMA3, and PSMC3, cells were lysed 15 h after H_2_O_2_ treatment. For HO-1, cells were lysed 18 h after H_2_O_2_ treatment. After centrifugation and supernatant collection, equal amounts of proteins were resolved by 10% SDS-PAGE and transferred to polyvinylidene fluoride membranes. After drying for 1 h at RT, the membranes were probed with p-Akt, Akt, Nrf2, p-CREB, PSMA3, PSMC3 (Cell Signaling Technology, Inc.), HO-1 (Enzo Life Sciences International Inc.), insulin-degrading enzyme (Abcam) and GAPDH (GeneTex, Inc.) antibodies overnight at 4°C. The washed membranes were then incubated with horseradish peroxidase-conjugated goat anti-rabbit secondary antibodies (Jackson ImmunoResearch Laboratories, Inc.). Immunoreactivity was visualized with an enhanced chemiluminescence substrate (Advansta, Inc.) and detected with the Amersham Imager 600 imaging system (GE Healthcare Life Sciences). The images were analyzed with Image Lab software (Bio-Rad Laboratories, Inc.).

### Statistical analysis

The results were presented as the mean ± standard deviation. Statistical analyses using one-way analysis of variance (ANOVA) followed by Tukey’s post-hot test were performed by OriginPro 2015 software (OriginLab). P-value < 0.05 was considered to indicate a statistically significant difference.

## Results

### In vitro-applied TC-HT

We applied the thermal cycle (TC) treatment to SH-SY5Y cells by a modified polymerase chain reaction (PCR) equipment (Fig 1C). Briefly, in our design, some protruding parts of the PCR machine and plastic well were milled so that the bottom of the well can touch the heat sink tightly. The basic TC settings were schemed as Fig 1A, where the temperature was elevated to the desired high temperature and sustained for a period of time followed by a cooling process at body temperature. It’s worth noting that the adjustment of TC parameters (temperature and time) is based on the needs and objectives of the experiments. In our experiments, the TC parameters were set to high temperatures of 42.5°C or 41.5°C for 15 min, then lowered to 37°C and lasted for 35 sec. This cycle was repeated 8 or 12 times. The HT group maintains the same high temperature setting throughout the treatment in a continuous manner without a break. To determine the actual temperatures of the cells, a needle thermocouple located in the bottom of the well was used to monitor the temperatures. Fig 1B represents the actual temperature measured by the thermocouple every 20 sec. The actual cycle temperature sensed by the cells was ~42-40.3°C for the setting of 42.5-37°C and ~40.9-39.4°C for 41.5-37°C setting, as shown in Fig 1B.

**Fig 1.**
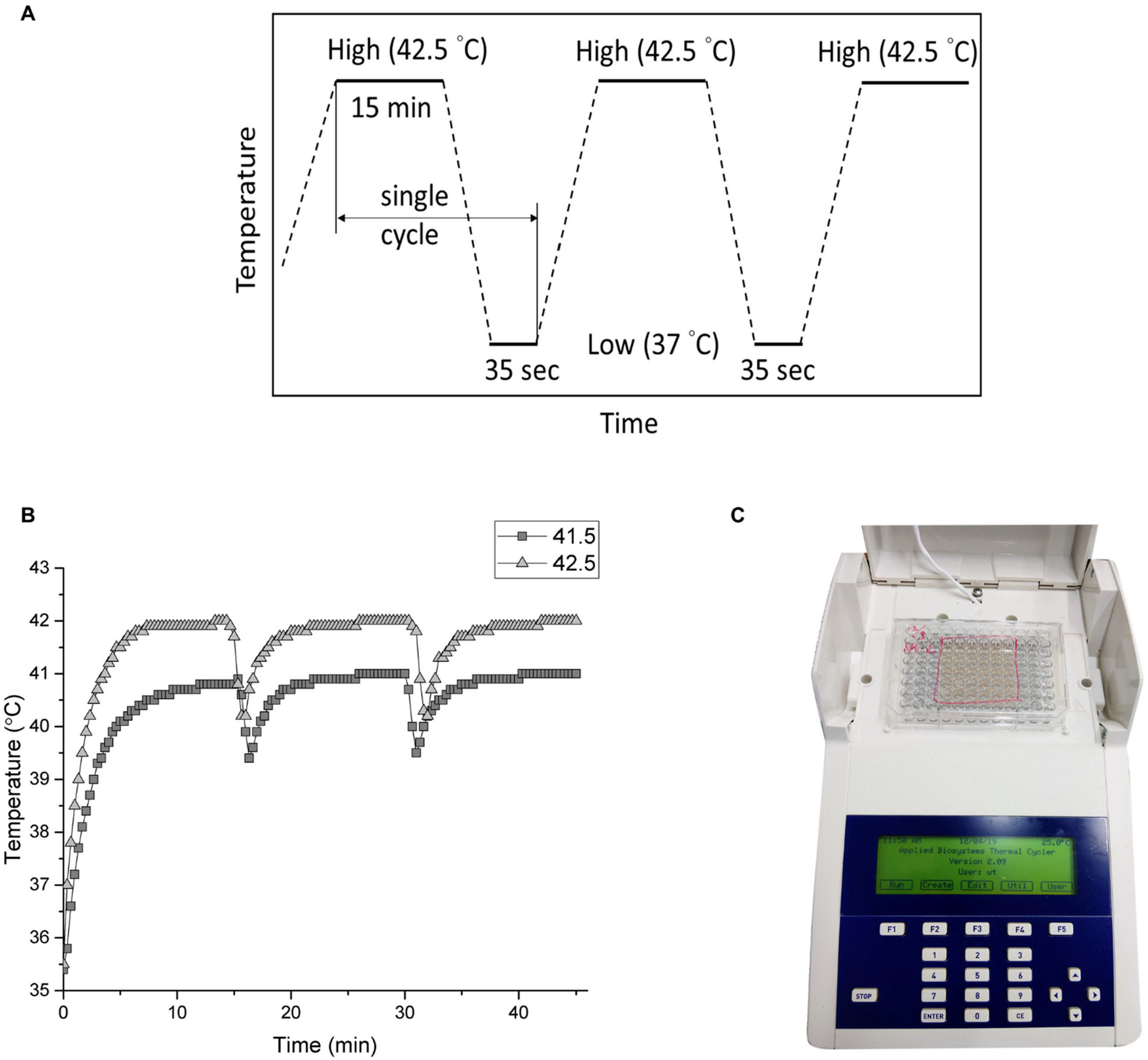
The setup of TC-HT. (A) Schematic representation of TC parameter settings. (B) Measurement of the cell temperature during the TC treatment by the thermocouple. (C) Image of the TC controller setup.

### Effect of TC-HT on H_2_O_2_ and Aβ-induced cytotoxicity in SH-SY5Y cells

We applied H_2_O_2_ to SH-SY5Y cells as an oxidative stress and the cell viability was assessed by MTT assay. As shown in Fig 2A, treatment of H_2_O_2_ on SH-SY5Y cells for 24 h induced significant decrease of cell viability in a dose-dependent manner. Treatment of SH-SY5Y cells with 450 µM H_2_O_2_ for 24 h reduced the cell viability to 52.9% compared to control cells, which was used in the following experiments. To investigate whether TC-HT or HT produced protection and increased cell survival against the oxidative stress induced by H_2_O_2_, the SH-SY5Y cells were pretreated with TC-HT or HT in the experiments. Two high temperature parameters (41.5°C or 42.5°C) were applied for 2 h or 3 h continuously in the HT group, and 8 or 12 cycles (for 15 min heating time per cycle) in the TC group to make the total thermal doses of the HT and TC groups equal. After the heat pretreatment, the treated SH-SY5Y cells were kept in the incubator at 37°C for 4 h. They were then challenged with H_2_O_2_ to generate the oxidative stress, and the cell viability was determined by MTT assay at 24 h after the H_2_O_2_ treatment. The results found that H_2_O_2_ treatment significantly reduced the viability of SH-SY5Y cells to 53.7% of the control value, and showed that the heat pretreatment conferred protective effect and increased the cell viability. At 41.5°C for 2 h and 8 cycles, HT and TC treatments increased the cell viability to 61.9% and 68.5% of the control value, respectively (Fig 2B). As we increased the high temperature setting to 42.5°C while maintaining the same heating time, TC pretreatment greatly increased the cell viability to 83% of the control value under H_2_O_2_ stress, while the HT group only exhibited a slight increase (Fig 2C). When we further increased the heating time to 3 h, the treatment of HT alone was cytotoxic to the cells, reducing the cell viability to 75.4%, and the viability was even lower after the H_2_O_2_ treatment (Fig 2C). Interestingly, the TC treatment with the same total thermal dose (12 cycles) had only a slight influence on the cell viability, and it still conferred significant neuroprotective effect to the SH-SY5Y cells under H_2_O_2_ stress.

**Fig 2.**
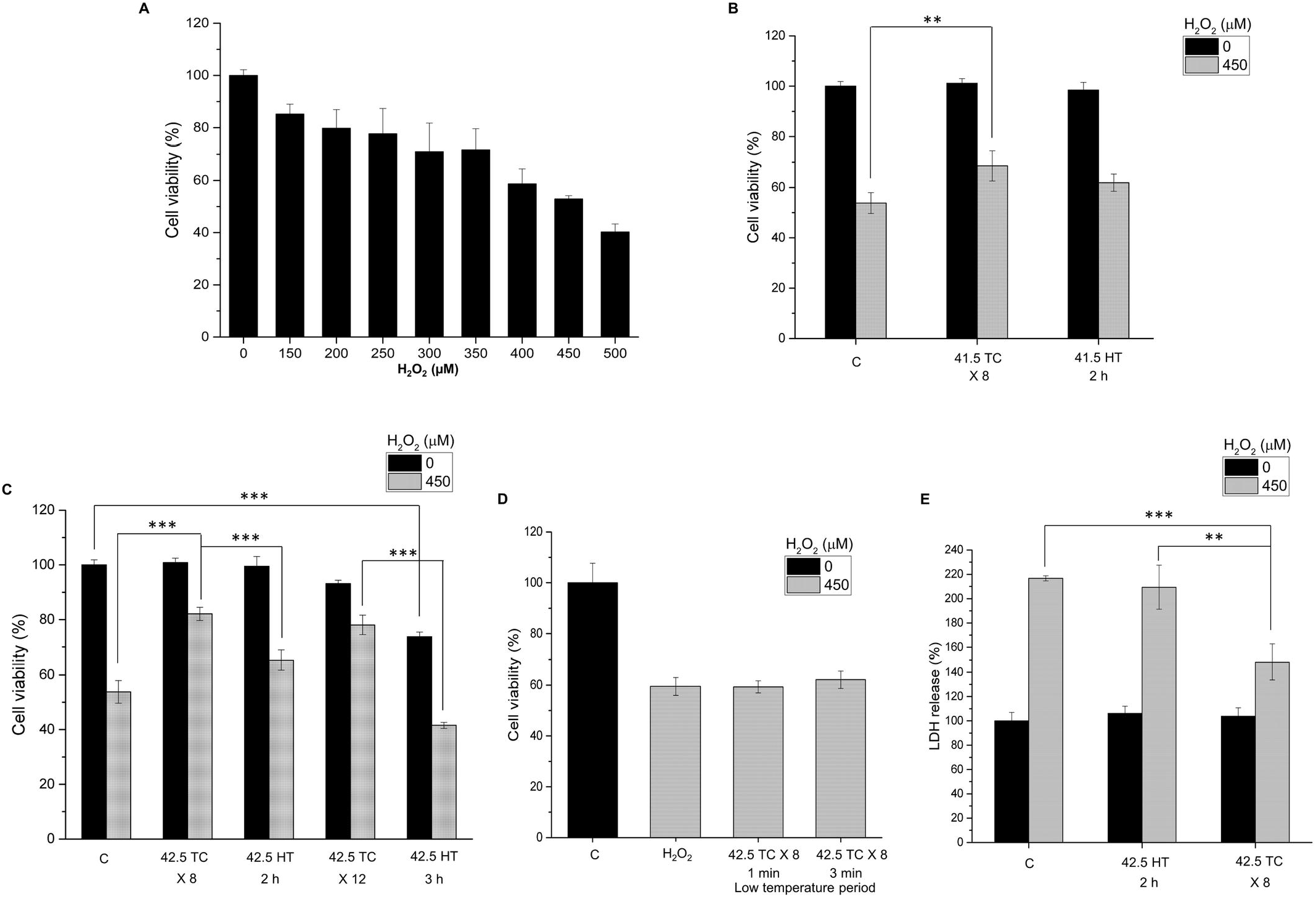
Effect of TC-HT on H_2_O_2_-induced cytotoxicity in SH-SY5Y cells. (A) Dose-response curve of SH-SY5Y cells treated with different concentrations of H_2_O_2_ for 24 h. (B) SH-SY5Y cells were pretreated at 41.5°C temperature setting and challenged with or without 450 µM H_2_O_2_. The cell viability was measured by MTT assay at 24 h after the H_2_O_2_ treatment. (C) SH-SY5Y cells were pretreated at 42.5°C temperature setting with different thermal dosages and challenged with or without 450 µM H_2_O_2_. The cell viability was measured by MTT assay at 24h after the H_2_O_2_ treatment. (D) Comparison of the neuroprotective effect under different low temperature period settings. (E) The LDH release was measured to confirm the neuroprotective effect of TC treatment. SH-SY5Y cells were pretreated at 42.5°C temperature setting and challenged with or without 450 µM H_2_O_2_. The LDH release was measured 24h after the H_2_O_2_ treatment. Data represent the mean ± standard deviation (n=3). ^***^P < 0.001 and ^**^P < 0.01.

In particular, the study examined the influence of the cooling process on the neuroprotective effect. As shown in Fig 2D, for low temperature period longer than 1 min, the TC pretreatment did not produce protective effect to the cells, indicating that the low temperature period is a critical parameter to be determined for effective neuroprotection. From molecular point of view, the neuroprotective effect would be discontinued as long as there were at least one or few critical signalling pathways being blocked as heating was halted, so we found that the low temperature period used for effective protection was shorter than the heating time in TC-HT treatment. On the other hand, the LDH release rate was measured and found to decrease significantly in the TC pretreatment group compared to the H_2_O_2_-treated group (Fig 2E), further confirming its protective role in maintaining the cell membrane integrity when cell was under oxidative stress. For AD disease model, Aβ treatment was used to reduce the viability of SH-SY5Y cells due to the cytotoxicity of the aggregated Aβ (Fig 3A). Our results showed that the TC treatment after Aβ administration greatly improved the viability of SH-SY5Y cells (Fig 3B) for nearly 25%, indicating the curative effect of TC in AD disease model in vitro. The most effective high temperature setting was 42.3°C for 8 cycles in the Aβ-treated cells. The actual high temperature sensed by the cells was about 41.7°C. Interestingly, the curative effect of TC pretreatment was less pronounced than that of TC post-treatment (treated after Aβ administration), indicating the time point of TC application has great impact on the curative effect. Moreover, there were no curative effects by both HT pretreatment and post-treatment on SH-SY5Y cells, whose viability levels were of no difference with that in the Aβ only group (Fig 3B), implying the toxicity of Aβ was not attenuated by heat treatment. Thus, the results confirmed the therapeutic effect of TC-HT in AD disease model in vitro. Furthermore, the light microscopy images showed that the integrity of the cells was destructed by Aβ, and the TC treatment caused the protective effect to retain the cell morphology, as shown in Fig 3C.

**Fig 3.**
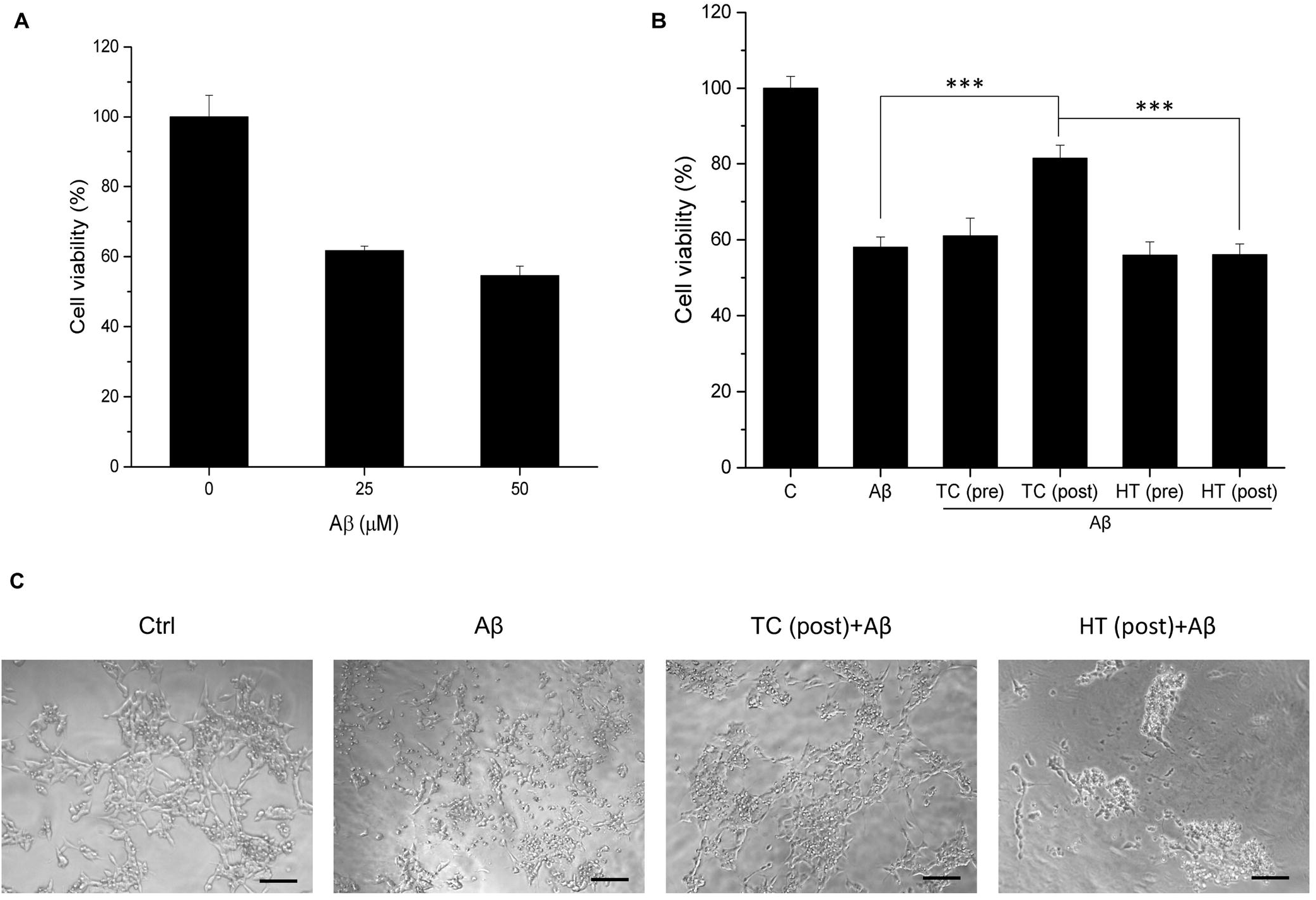
Effect of TC-HT on A β-induced cytotoxicity in SH-SY5Y cells. (A) The cell viability of SH-SY5Y cells treated with 25 or 50 µM Aβ for 4 days. (B) The TC or HT pretreatment and post-treatment at 42.5°C temperature setting were applied to the cells before or after 50 µM Aβ administration, and the cell viability was measured by MTT assay 4 days after treatment. The TC or HT treatment was applied 4 h before the Aβ administration for the pretreatment group and 1 h after Aβ administration for the post-treatment group. (C) Representative light microscopy images of SH-SY5Y cells after treatment. The integrity of the cells was destructed by Aβ, and the TC post-treatment caused the protective effect and retained the cell morphology. Scale bar = 100 µm. Data represent the mean ± standard deviation (n=3). ^***^P < 0.001.

To conclude the result of the H_2_O_2_-induced oxidative stress on SH-SY5Y cells, the protective effect of TC pretreatment was more pronounced than the HT preconditioning. And the TC pretreatment at 42.5°C produced much superior protection effect than at 41.5°C. Noteworthily, the HT pretreatment at 42.5°C for 3 h decreased the cell viability which indicates that the continuous heating may induce cell damage. The protective effect of TC was most remarkable at ~42.5°C for 8 cycles. Therefore, we chose this parameter to evaluate the possible protective mechanism of TC-HT in the following experiments.

### TC-HT attenuates H_2_O_2_-induced ROS generation

The onset and progression of AD has been reported to be associated with ROS. We further studied whether the H_2_O_2_-induced intracellular ROS production could be attenuated by the TC pretreatment. The intracellular ROS levels were detected using the fluorescent dye DHE (Fig 4A). Results showed that after exposure to 450 µM H_2_O_2_ for 24 h, ROS level was increased to 160% of the control value. When the SH-SY5Y cells were pretreated with TC-HT, the intracellular ROS level was significantly ameliorated (Fig 4B). Compared to the TC group, the ROS level was not changed in the HT group.

**Fig 4.**
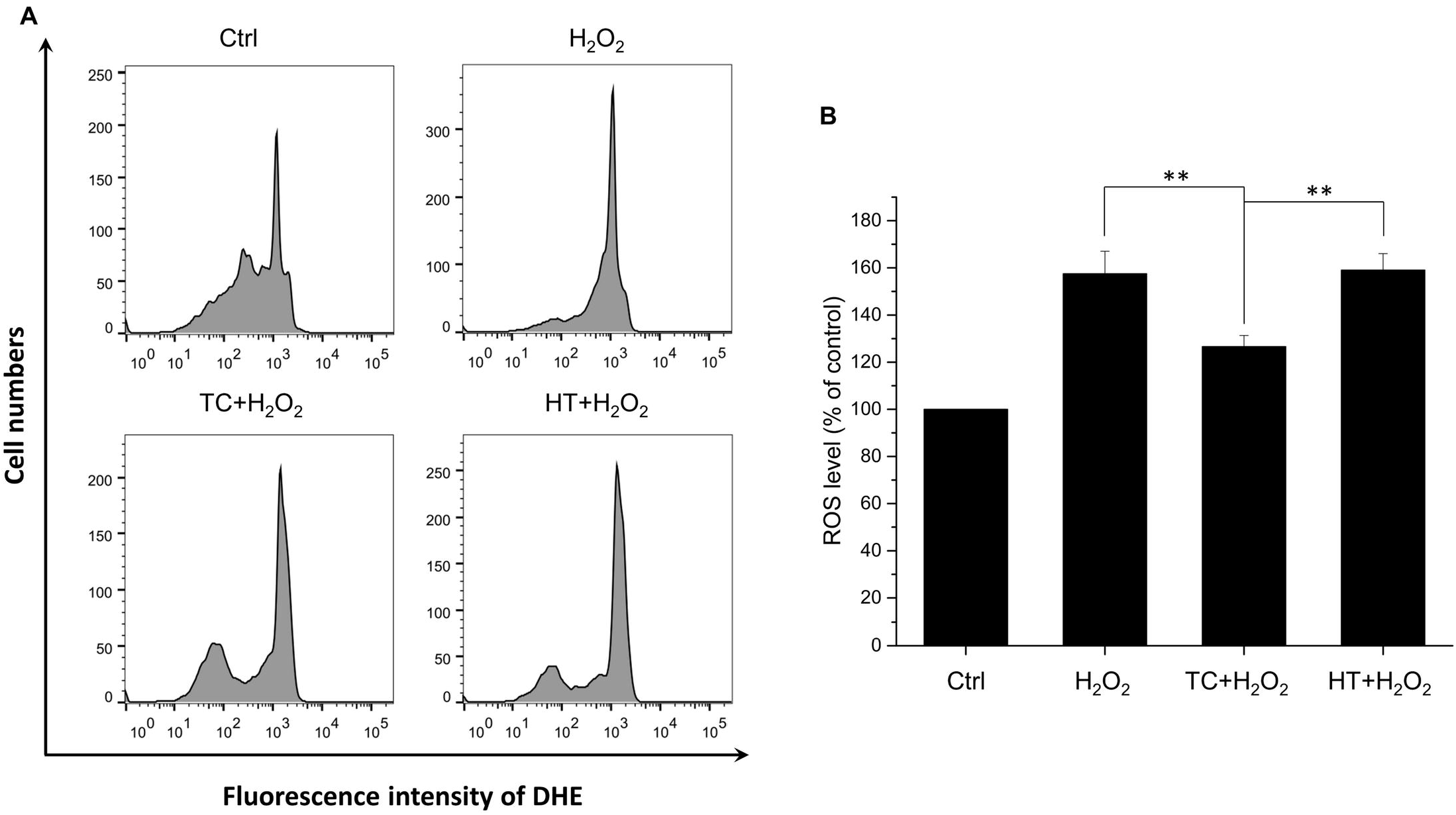
Effect of TC-HT on H_2_O_2_-induced ROS generation in SH-SY5Y cells. (A) ROS level was measured 24 h after the H_2_O_2_ treatment by flow cytometry with DHE fluorescent dye. (B) Quantification of the ROS levels after H_2_O_2_, TC+H_2_O_2_, or HT+H_2_O_2_ treatment. Data represent the mean ± standard deviation (n=3). ^**^P < 0.01.

### TC-HT attenuates MMP loss in H_2_O_2_-treated cells

Many lines of evidence have suggested that mitochondria contribute to ageing-related neurodegenerative diseases through the accumulation of net cytosolic ROS production [21]. The levels of MMP are kept relatively stable in healthy cells. Apoptotic signals are usually initiated by the disruption of normal mitochondrial function, especially the collapse of MMP. To study the mechanism why H_2_O_2_-induced apoptosis is alleviated by TC-HT pretreatment, the MMP is further analyzed by the flow cytometry with DiOC_6_(3) fluorescent dye in the study. As shown in Fig 5A, the oxidative stress induced by H_2_O_2_ caused mitochondrial dysfunction, decreasing the MMP and therefore decreased the intensity of the DiOC_6_(3) fluorescent signal. From the quantification results (Fig 5B), it was found that the percentage of cells with decreased MMP drastically increased after the H_2_O_2_ treatment. In contrast to the group of H_2_O_2_ alone, the TC-HT pretreatment noticeably suppressed the H_2_O_2_-induced dissipation of MMP, so the cells with decreased MMP reduced significantly, which was attributed to the preservation of the mitochondrial function and thus caused the blockage of the apoptotic signaling transduction.

**Fig 5.**
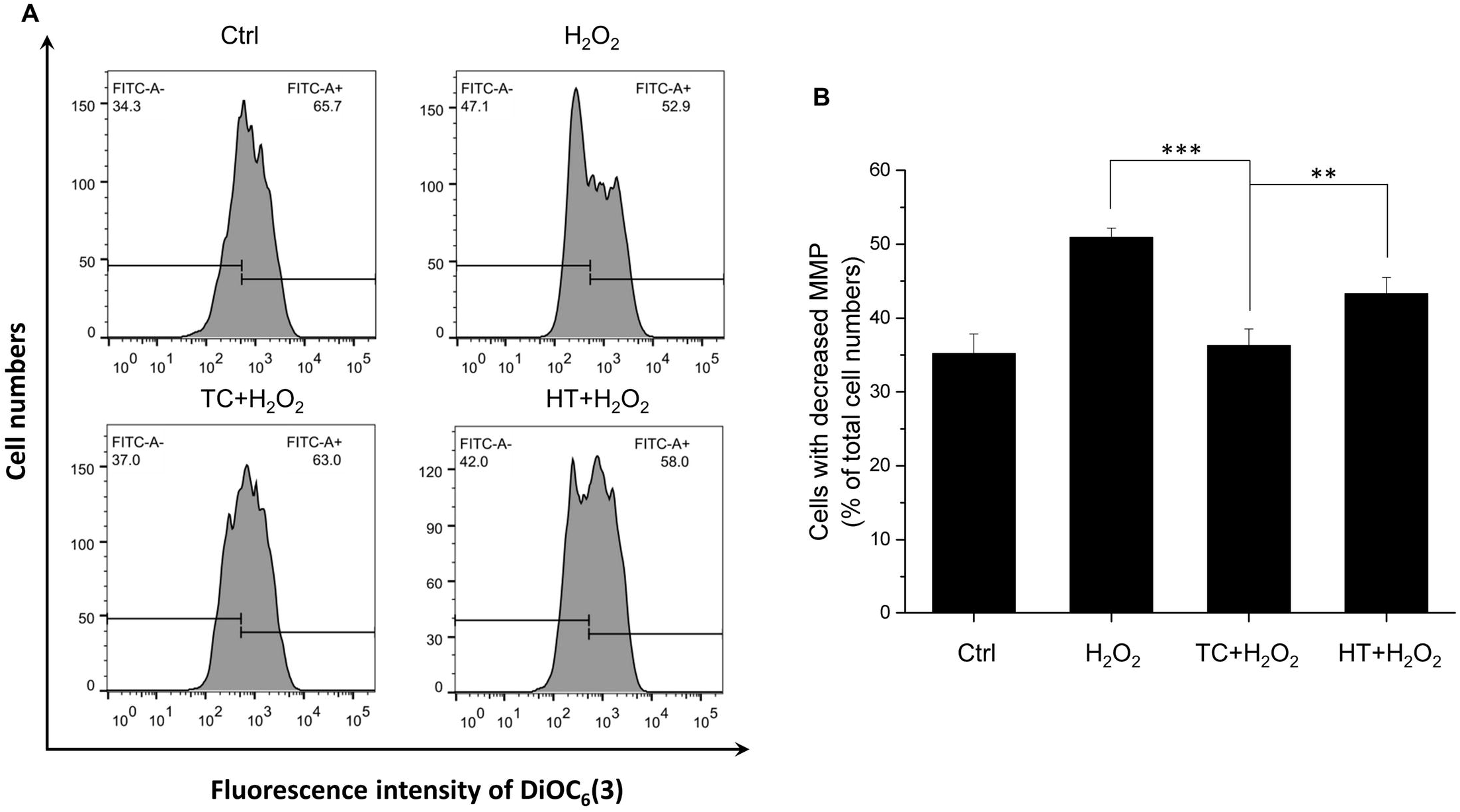
Effect of TC-HT on H_2_O_2_-induced MMP reduction in SH-SY5Y cells. (A) MMP was analyzed 24 h after the H_2_O_2_ treatment by flow cytometry with DiOC_6_(3) fluorescent dye. (B) Quantification of the cells with decreased MMP after H_2_O_2_, TC+H_2_O_2_, or HT+H_2_O_2_ treatment. Data represent the mean ± standard deviation (n=3). ^***^P < 0.001 and ^**^P < 0.01.

### Effect of heat treatment on heat shock protein (HSP) expressions in SH-SY5Y cells

It has been well known that HT triggers expression of HSPs so that cells are able to protect themselves from stress [22, 23]. To investigate whether the protective effect of TC-HT was related to the HSPs, we examined the protein expressions of HSP70 and HSP105, both of which were reported to have protective effects. As shown in Fig 6, both HSP expressions were enhanced significantly after HT and TC treatments. The activation levels of TC, however, was only slightly higher than HT without significant difference. This phenomenon points out that although TC treatment indeed protects SH-SY5Y cells more from the oxidative stress of H_2_O_2_, the main mechanism could be apart from the HSP70 or HSP105. Therefore, other heat-activated stress response proteins should be considered in the experiments.

**Fig 6.**
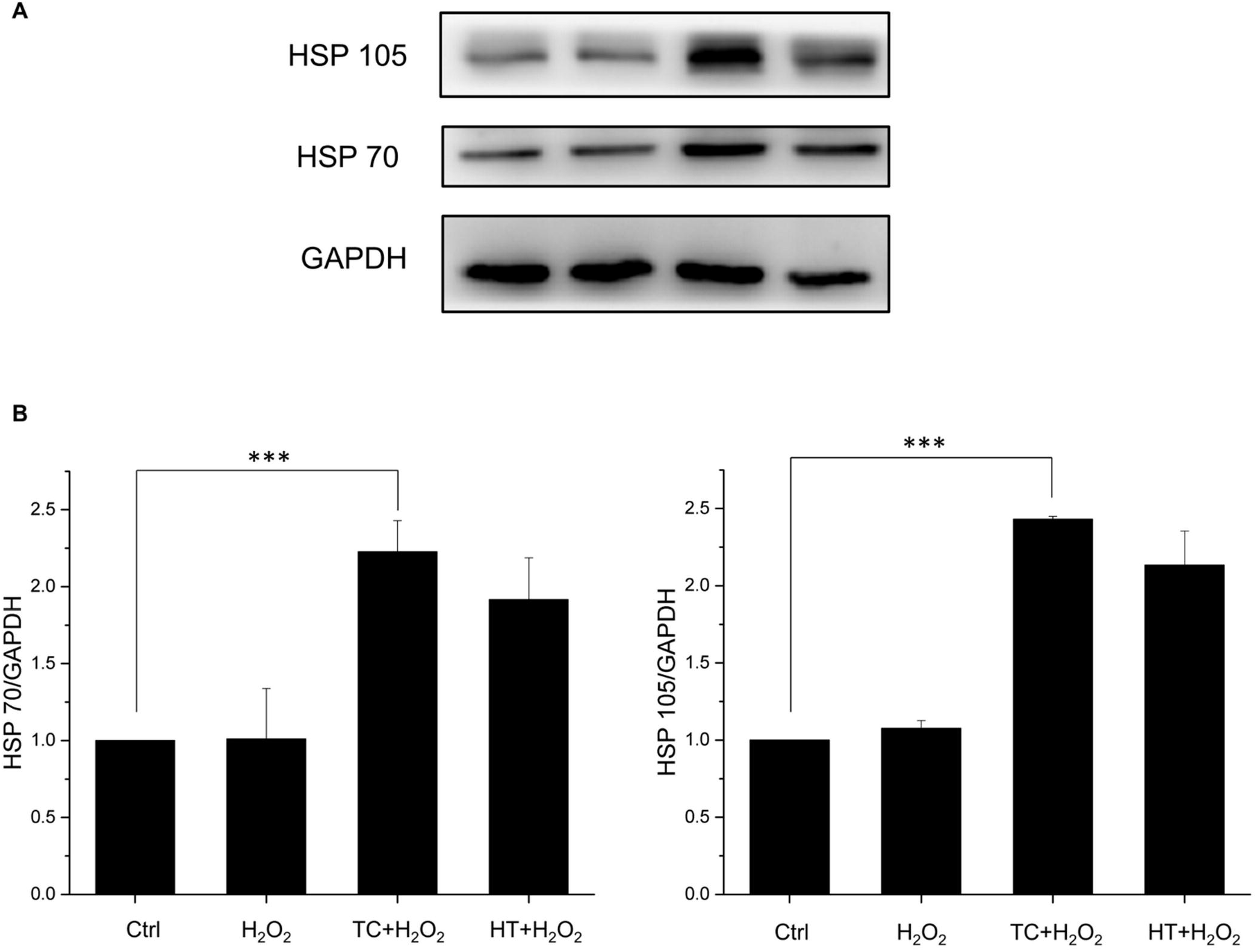
Effect of TC-HT on expressions of HSPs in SH-SY5Y cells. (A) Cells were lysed 16 h after H_2_O_2_ treatment and western blot analyses of HSP70 and HSP105 expressions were performed. (B) Quantification of HSP70 and HSP105 expressions after H_2_O_2_, TC+H_2_O_2_, or HT+H_2_O_2_ treatment. The expression levels were normalized to GAPDH. Data represent the mean ± standard deviation (n=3). ^***^P < 0.001.

**Fig 7.**
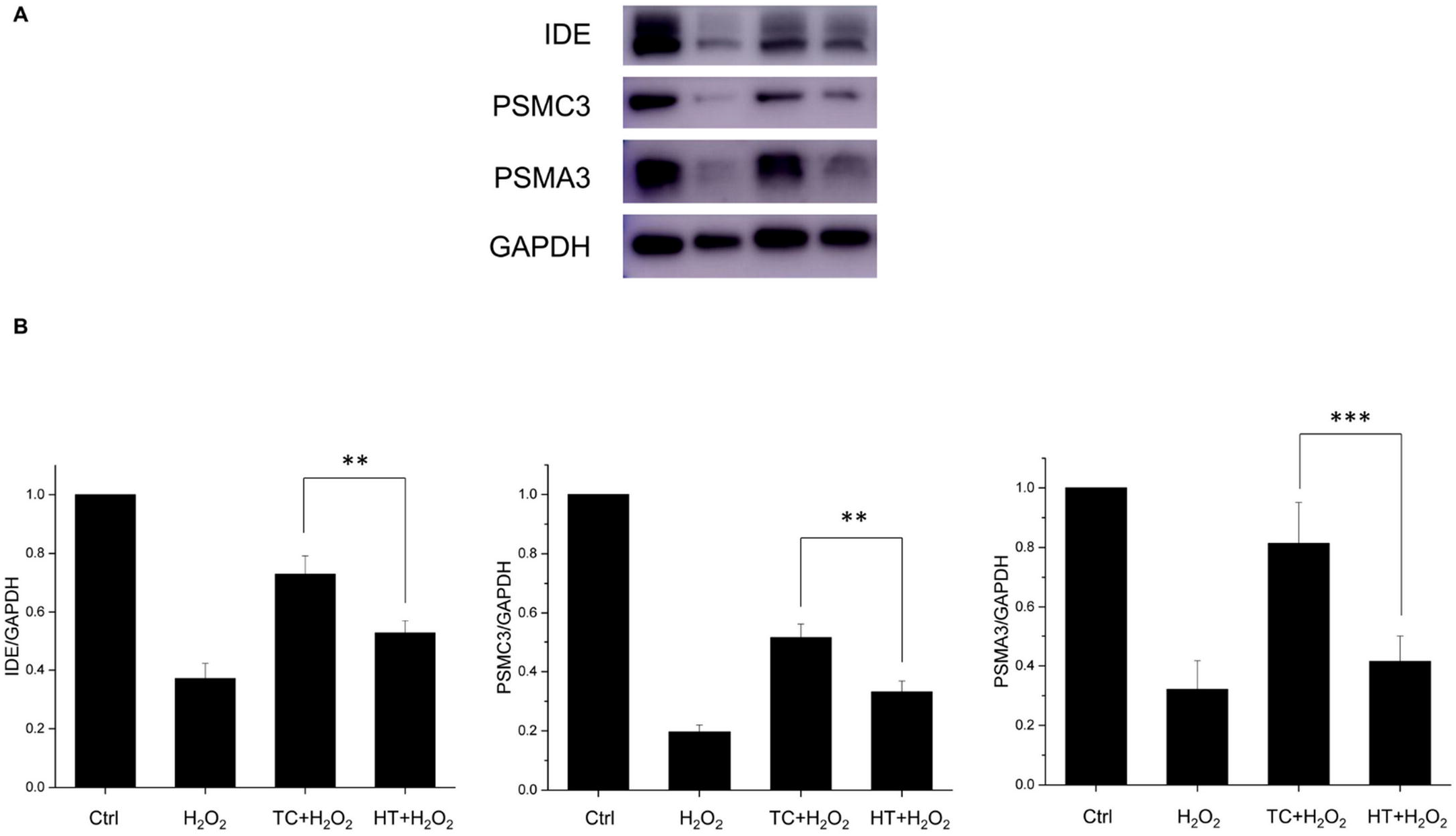
Effect of TC-HT on expressions of IDE and proteasome in SH-SY5Y cells. (A) Cells were lysed 15 h after H_2_O_2_ treatment and western blot analyses of IDE and proteasome subunits (PSMC3 and PSMA3) expressions were performed. (B) Quantification of IDE, PSMC3, and PSMA3 expressions after H_2_O_2_, TC+H_2_O_2_, or HT+H_2_O_2_ treatment. The expression levels were normalized to GAPDH. Data represent the mean ± standard deviation (n=3). ^***^P < 0.001 and ^**^P < 0.01.

### Effect of heat treatment on insulin-degrading enzyme and proteasome expressions in SH-SY5Y cells

Insulin-degrading enzyme (IDE) is a highly conserved zinc metallopeptidase which was originally identified as the key enzyme for the degradation of insulin. Further studies unraveled its ability to degrade several other polypeptides, including Aβ [24]. Recent studies demonstrated that IDE was up-regulated in a HSP-like fashion [25]. In this study, we also examine the expression levels of IDE under the TC or HT treatment. The results showed that H_2_O_2_ significantly decreased the expression level of IDE and the heat treatment by TC remarkably recovered its expression level while HT had notably weaker ability to regain the normal condition. Another important protein quality control system which could degrade the Aβ peptide is the ubiquitin-proteasome system (UPS). Regulation of Aβ production by the UPS may have a role in AD pathogenesis and therefore the UPS is regarded as potential therapeutic target for AD [26]. The proteasome comprises a proteolytic 20S core particle capped by either one or two 19S regulatory particles. In this study, the protein expression levels of 20S (PSMA3) and 19S (PSMC3) subunits were examined to verify the effect of thermal treatment on proteasome expression. The results showed that both expressions of PSMC3 and PSMA3 were weakened by H_2_O_2_ significantly. For the thermal treatment by TC, the levels of PSMC3 and PSMA3 expressions were greatly recovered especially in the PSMA3 level (return to 80% of the control value). For the HT treatment group, the degree of recovery was much lower than TC treatment group in both PSMC3 and PSMA3 expressions, while it had no significant difference compared to the H_2_O_2_-treated group in PSMA3 protein expressions.

### Effect of heat treatment on Akt-Nrf2/CREB signalling pathways in H_2_O_2_-treated SH-SY5Y cells

We then examined the signalling pathways that could participate in the neuroprotective mechanism of TC-HT. The Akt pathway is highly conserved in all cells of higher eukaryotes and plays pivotal roles in cell survival and growth. Once Akt is activated by phosphorylation, it positively regulates some transcription factors to allow expression of prosurvival proteins. Therefore, in our study, we examined the phosphorylation of Akt and the downstream transcription factors including cAMP response element binding protein (CREB) and nuclear factor erythroid 2-related factor 2 (Nrf2) by western blot analysis. As shown in Fig 8, the expression of phosphorylated Akt (p-Akt) was increased in H_2_O_2_-treated SH-SY5Y cells compared to the untreated cells. The increased p-Akt level could be due to the stress response induced by the oxidative stress of H_2_O_2_. Heat pretreatment further increased the p-Akt levels in H_2_O_2_-treated SH-SY5Y cells, and the TC group had significantly higher level of p-Akt than the HT group. We also examined the expression levels of p-CREB and Nrf2. CREB is an important transcription factor that also triggers expression of many prosurvival proteins, including Bcl-2 and brain derived neurotrophic factor (BDNF) [27]. When CREB is phosphorylated, it starts to regulate downstream gene expression. Besides, Nrf2 is also a key transcription factor that regulates expression of antioxidant proteins, such as heme oxygenase-1 (HO-1) [28]. As shown in Fig 8, H_2_O_2_ treatment slightly influenced Nrf2 and p-CREB protein expression levels. In the heat-treated groups, the TC treatment significantly increased the expressions of Nrf2 and p-CREB, while the HT treatment increased Nrf2 expression with less significant level than TC treatment but in turn reduced the expression level of p-CREB. The HO-1 protein is a heme-containing HSP which is crucial for neuroprotection against oxidative stress. The result showed that the heat treatment significantly enhanced the expression levels of HO-1 in TC and HT groups. Particularly, it is worth noting that the TC treatment had a significant higher level of HO-1 than HT, which could account for the antioxidant ability of the TC treatment in the ROS experiment.

**Fig 8.**
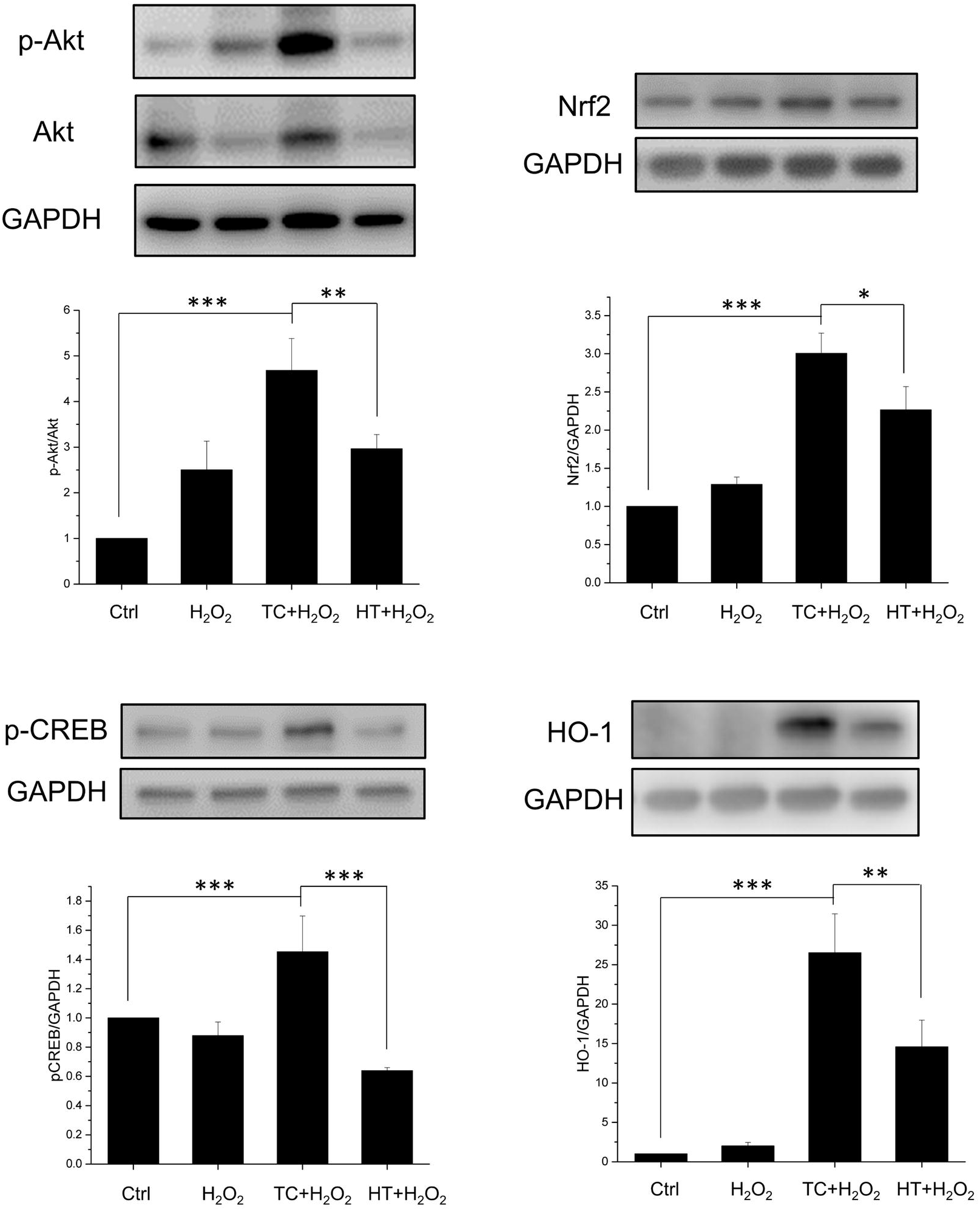
Effect of TC-HT on Akt/Nrf2 and Akt/CREB signalling pathways and related protein expressions. Cells were lysed 15 h after H_2_O_2_ treatment for p-Akt, Akt, Nrf2, p-CREB proteins, and lysed 18 h after H_2_O_2_ treatment for HO-1 protein, and western blot analyses of p-Akt, Akt, Nrf2, p-CREB, and HO-1 proteins were performed. Quantification of p-Akt, Nrf2, p-CREB, HO-1 expressions after H_2_O_2_, TC+H_2_O_2_, or HT+H_2_O_2_ treatment. The expression level of p-Akt was normalized to total Akt while other proteins were normalized to GAPDH. Data represent the mean ± standard deviation (n=3). ^***^P < 0.001, ^**^P < 0.01 and ^*^P < 0.05.

### Effect of LY294002 on the TC-HT activated Akt pathway and protective effect

In order to examine if the activation of the Akt pathway was involved in the neuroprotective effect of TC treatment, we inhibited the phosphoinositide-3-kinase (PI3K), the upstream kinase of Akt, by the inhibitor LY294002 [29] in the study. In Fig 9A, we found that H_2_O_2_ treatment reduced the viability of SH-SY5Y cells to 58% of the control value, and TC treatment could greatly increase the cell viability to ~86% of the control value under H_2_O_2_ stress. In the presence of PI3K inhibitor, the p-Akt level was indeed reversed and the neuroprotective effect of TC treatment against H_2_O_2_-induced cell apoptosis was abrogated (Fig 9A and 9B), representing that the Akt pathway was involved to mediate the neuroprotective effect of TC treatment against H_2_O_2_ toxicity. For the downstream proteins of Akt, inhibition of the PI3K/Akt pathway was found to significantly reduce the induction of Nrf2 by TC treatment, indicating that the transcription factor Nrf2 was indeed activated by the Akt pathway. Collectively, these data point out that TC treatment may exhibit neuroprotective effect via activation of the Akt-Nrf2/CREB signalling pathways in the H_2_O_2_-treated SH-SY5Y neuron cells.

**Fig 9.**
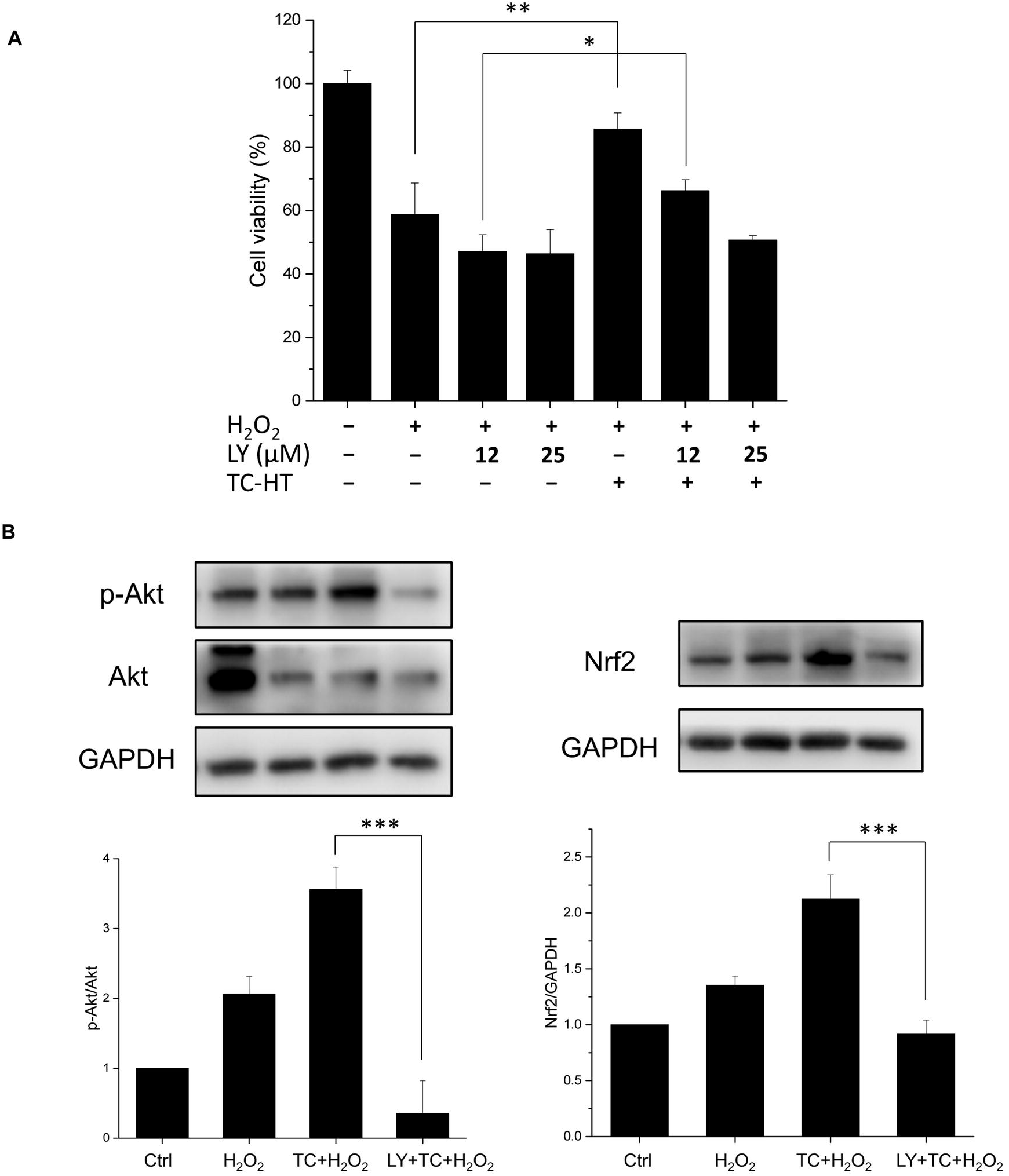
Effect of LY294002 on the neuroprotective effect of TC-HT and related protein expressions. (A) TC treatment conferred neuroprotective effect and significantly increased the cell viability of SH-SY5Y neuron cells under the H_2_O_2_ stress. The neuroprotective effect of TC treatment was abrogated by addition of the PI3K inhibitor LY294002 in a dose-dependent manner. (B) Cells were lysed 15 h after H_2_O_2_ or 25 µM LY294002 treatment, and western blot analyses of p-Akt and Nrf2 proteins were performed. The inhibitor LY294002 reversed the activated levels of p-Akt and Nrf2 induced by TC treatment. The expression level of p-Akt and Nrf2 was normalized to total Akt and GAPDH, respectively. We used the abbreviation “LY” to represent the PI3K inhibitor LY294002 in the figure. Data represent the mean ± standard deviation (n=3). ^***^P < 0.001, ^**^P < 0.01 and ^*^P < 0.05.

## Discussion

The study focused on investigating the neuroprotective effect of TC-HT on SH-SY5Y cells. It showed that subjection of the cells to heat at high and low temperatures alternately could produce significant protective effect against H_2_O_2_ and Aβ-induced apoptosis in SH-SY5Y human neural cells. The western blot results showed that TC-HT activated the PI3K/Akt signalling pathway and mediated the activation of Nrf2 and CREB proteins.

A human derived cell line SH-SY5Y has been extensively employed as general in vitro model in evaluating neuronal damage or neurodegenerative diseases, such as AD and PD [30, 31]. Therefore, the study used SH-SY5Y cells in the experiments to determine the protective effect of TC-HT against H_2_O_2_ and Aβ-induced cytotoxicity. Although the exact molecular mechanism of neurodegeneration pathogenesis is still not clear, a common feature of these diseases is oxidative stress [32]. Many researchers believe that oxidative stress may play a critical role in the etiology and cause the exacerbation of disease progression. Cellular ROS are generated by both extrinsic and intrinsic sources, with the former including ultraviolet, drugs, and environmental toxins. Under normal physiological conditions, ROS generated intrinsically from mitochondria and other enzymes are maintained at appropriate levels by endogenous antioxidants. However, when mitochondria suffers from decreased antioxidant defense or cell inflammation due to damage, excess ROS production may occur. Since neuron cells are especially vulnerable to oxidative damage due to their high oxygen demand and high polyunsaturated fatty acid contents in membranes [33], the imbalance between ROS production and removal may cause neuron damage or degeneration. A key cellular and molecular mechanism of several neurodegenerative diseases, such as AD, PD, Huntington’s disease (HD), amyotrophic lateral sclerosis (ALS), is the accumulation of misfolded aggregation proteins in brain [34]. The deposition of these misfolded protein aggregates can cause inflammation in brain, inducing significant ROS production and leading to synaptic dysfunction, neuronal apoptosis, and brain degeneration [35]. The composition of protein aggregates is different among various neurodegenerative diseases. For AD, there are two main kinds of protein aggregates, namely extracellular amyloid plaques deriving from amyloid precursor protein and intracellular microtubule-associated protein tau, known as neurofibrillary tangles [36]. With the complex pathogeneses of Aβ or tau proteins still remaining largely unknown, several studies have proven that the accumulation of Aβ increased oxidative stress and led to mitochondrial dysfunction and DNA/RNA oxidation [37]. Matsuoka et al. showed that AD transgenic mouse with Aβ accumulation can cause an increase in H_2_O_2_ level, suggesting that Aβ may enhance oxidative stress in AD [38]. The oxidative stress can boost the hyper-phosphorylation of tau protein and in turn facilitate the aggregation of Aβ [39]. Studies have also pointed out that overexpressed tau proteins make the cells more vulnerable to oxidative stress [40]. Therefore, efforts have been underway to find therapeutic strategies that can protect against oxidative stress and alleviate the symptoms of neurodegenerative diseases. Among numerous ROS species, H_2_O_2_ can be considered to be one of the most important ROS molecules involved in the progression of AD. Many studies, therefore, used H_2_O_2_ to induce oxidative stress in vitro, as a model in the investigation of the neuroprotective or neurodamage mechanisms [41–43]. The study also found that H_2_O_2_ increased the ROS level in SY-SY5Y cells and caused the cell death in a dose-dependent manner. Additionally, Aβ was used to further investigate the neuroprotective effect of TC-HT in the AD disease model. Both MTT and LDH release assays were employed in assessing the neuroprotective effect of TC-HT. The results showed that TC-HT increased the cell viability in the H_2_O_2_ and Aβ-treated neural cells. The detection of ROS level showed that one possible mechanism of neuroprotective action of TC-HT exhibited the antioxidant property, reducing the intracellular ROS level, perhaps due to the increase of antioxidant enzymes in SH-SY5Y cells, induced by TC-HT. In the western blot results, the study proved that such effect was through activation of Akt/Nrf2 pathway.

The Akt signaling pathway is a key regulator in the survival, proliferation, and migration of cells, in response to extracellular signals [44]. Phosphorylation of Akt further activates a set of downstream transcription factors, including Nrf2 and CREB, which are considered to be mediators of neuroprotection by increasing the expression of many antioxidant and prosurvival enzymes under oxidative stress. Nrf2 is one of the most important transcription factors that regulate the expression of various antioxidant proteins and maintain the redox state in mammalian cells [45, 46]. Among various antioxidant proteins, HO-1 has been proven to be effective in promoting antioxidant production and protection against neuronal injury [28]. The study found that the physical stimulation, TC-HT, could activate the Akt/Nrf2 pathway and upregulate the expression of the antioxidant protein HO-1, thereby reducing the ROS generation by H_2_O_2_. Compared with the continuous heating of HT group, TC-HT performed significantly better in activating Akt/Nrf2, HO-1 expression, and ROS reduction, in agreement with the cell viability assay.

Another possible mechanism for neuroprotection of TC-HT may be related to the activation of Akt/CREB pathway. CREB is the cellular transcription factor activated by phosphorylation from Akt or other kinases, which triggers expression of neurotrophins and antiapoptotic proteins. The critical role of CREB in neuronal plasticity and long-term memory has also been well-documented [47]. In addition to the neuroprotective effect of CREB by upregulating neurotrophins and antiapoptotic proteins, recent studies have discovered that CREB also protects neurons via the antioxidant pathways [27]. The downstream antioxidant genes include HO-1 and manganese superoxide dismutase (MnSOD). Besides, CREB downregulation is involved in the pathology of AD [48], and thus increasing the expression of CREB has been considered a potential therapeutic strategy for AD [49]. In some preconditioning experiments, CREB seemed to play a key role in the preconditioning response which protected the cells from the subsequent stress [50]. In the study, it was observed that TC-HT pretreatment significantly upregulated the activation of CREB protein, compared with the control group in SH-SY5Y cells. The result further found that HT significantly decreased the expression level of p-CREB compared with the control cells. The decreased level of p-CREB in HT group indicates that HT may exert too much heat stress and thus hinder the neuroprotective effect or even cause damage to the cells. Taken together, the result points out that the neuroprotective effect of TC-HT could be attributed to the function of antioxidant protein HO-1 or other prosurvival proteins via activation of Nrf2 and CREB through PI3K/Akt pathway (Fig 10). Further studies are necessary to identify the specific role of other downstream neurotrophins and antiapoptotic or antioxidant proteins in the neuroprotection effect.

**Fig 10.**
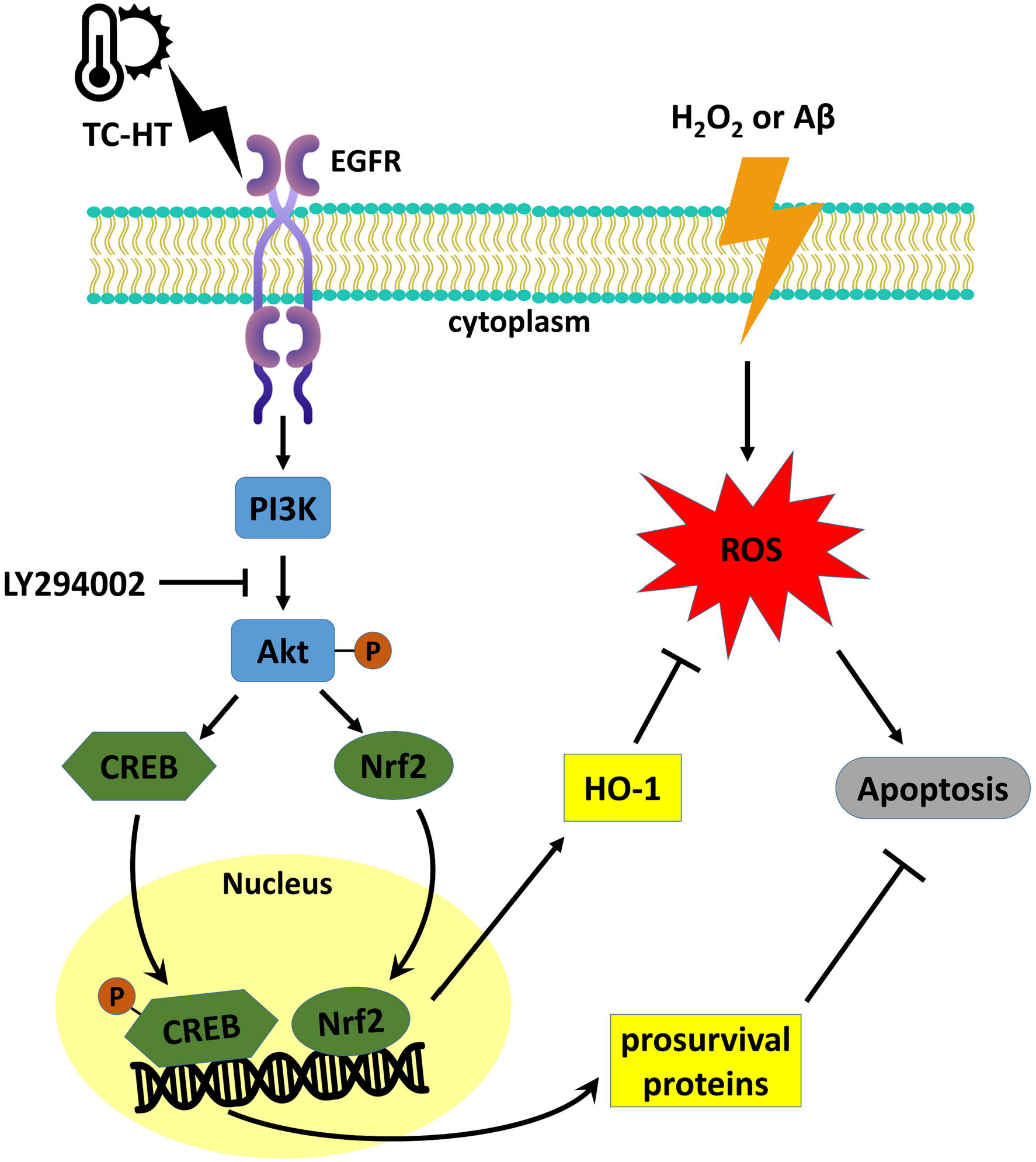
The proposed mechanisms for the protective effect of TC-HT against H_2_O_2_ or Aβ-induced neural injury. The epidermal growth factor receptor (EGFR) in the cell membrane could be the upstream receptor sensing extracellular TC-HT stimulation and transmit the signal to activate the PI3K/Akt pathway, which induces Nrf2 and CREB activations. The transcription factors entering the nucleus further enhance the expressions of HO-1 and other prosurvival proteins which decrease the ROS level and inhibit the apoptosis signal.

Under stress amid a harsh environment or caused by external stimuli, cells or whole organisms would trigger self-defense systems in response [51]. For example, fever induced by viral or bacterial infections is a useful defense mechanism capable of regulating the body temperature, as higher body temperature strengthens immune cells and increases their ability to kill bacteria and viruses. It also inhibits the growth of bacteria and viruses, thereby attaining anti-inflammatory effects. Depending on the intensity and duration of stress, cells would activate cell repair or stress response to adapt to the new environment or cause their death [52]. The stress response involves the synthesis of some highly conserved proteins with protective function for cells [53]. For the thermal stress induced when cells or whole organisms are exposed to elevated temperatures, they would respond by synthesizing an evolutionary highly conserved family of proteins, known as HSPs [54]. The HS response is a universal phenomenon, which appears in every tested organism. HSPs are named and classified, according to their molecular weight. HSP70 (70 kilodaltons in size) is a typical group of HSPs, which functions a molecular chaperone, to prevent the misfolding of nascent polypeptide chains and facilitate refolding of misfolded proteins [22]. Many studies have indicated that HSP70 sub-family is essential in protecting organisms from various stresses [55]. The study also examined the effect of TC-HT and HT on the expression of HSP70 and HSP105 [23], both of which have been reported to have protective effects. The results showed that HT and TC-HT treatments enhanced the expression of both HSPs significantly. However, the activation level of TC-HT was only slightly higher than HT group, which could not explain the superior protective effect of TC-HT compared to HT. Another novel HS-like protein, IDE, was identified as the major Aβ degrading enzyme, and its protein expression level was found to correlate negatively with the AD pathology [24]. Previous study had shown that IDE expression was stress-inducible [25]. In the study, it was found that the TC treatment greatly recovered the H_2_O_2_-induced IDE reduction, and the enhanced level was much higher than HT treatment. Therefore, IDE could be one of the proteins participating in the neuroprotective effect under Aβ administration.

The stress response enables cells to survive the stress-induced damage. Use of cells’ self-defense ability to adapt the stress environment or manipulation of the stress response to increase the survival rate may be valuable in regenerative medicine or disease curing. In fact, there have been examples involving preconditioning of cells with non-lethal stress to minimize cell damage and improve the transplantation outcome [56]. While there have been many experiments on thermotolerance and cell survival [57], none have demonstrated the stress response in neuroprotection and curation to the best of our knowledge. Besides, the challenge for the manipulation of cellular stress response is how to fine-tune stress intensity and duration to suit specific needs and prevent the stress-induced cell damage. The present study demonstrated a delicate and efficient way to activate the stress response, leading to protection and curation against H_2_O_2_ and Aβ-induced cytotoxicity on SH-SY5Y cells. The thermal dosage was fine-tuned in a heat-and-cold cycling process, to maximize the protective effect and minimize the heat stress-induced cell damage. Although traditional HS can activate HSPs which may provide resistance to insults, such as hyperthermia, the continuous exposure may cause cell damage or even death. Previous studies have shown that the threshold for thermal damage is different among various tissue cells, which have different levels of thermal sensibility [58]. For example, the skin cells can withstand much higher thermal dosage (47°C for 20 min) than neural cells, because the neural cells are more vulnerable. As the most thermally sensitive tissue, brain can be damaged by low thermal dosage [58]. To attain optimal thermal dosage for various tissues, the heating process should be divided into several parts, which is the basic architecture of TC-HT.

In this study, the work presented an efficient and guaranteed way of controlling the thermal dosage applied to cells. The novelty of the TC-HT strategy is periodically interspersing the short period of cooling process in the continuous heating HT treatment. The body temperature (37°C) was selected as the low temperature setting which mimics the passive cooling process in the human body. Interestingly, our data demonstrated that the application time of the low temperature period is a critical parameter to determine the efficacy of neuroprotection for TC-HT. For too short cooling period, the accumulation of the thermal dosage will cause damage to neural cells like HT. On the other side, it was found that the TC-HT did not produce protective effect to the cells when low temperature period was longer than 1 min (Fig 2D). It may be due to the reason that the application of too long cooling period (> 1 min) could interrupt the transmission of biochemical signals and protein expressions stimulated by thermal stress, thus failing to achieve the neuroprotective effect. The results revealed that 35 sec cooling process at 37°C after 15 min heating at 42.5°C for 8 cycles produces the optimum protective effect to SH-SY5Y cells. This protective effect employing TC-HT was significantly much better than the continuous HT, which was due to the activation of other thermal stress-associated proteins apart from HSPs and the prevention of heat damage. From the LY294002 experiment, it was confirmed that the superior protective effect resulted partially from the activation of Akt pathway. One fascinating phenomenon is the antioxidant effect induced by TC-HT, which could be attributed to the increased antioxidant protein HO-1. Besides, the Aβ-induced cytotoxicity was also rescued in TC-HT treated cells, which points out the possibility that some stress-induced proteins might have the ability to eliminate or refold the aggregated protein or ease the symptom at least. The chaperone or proteasome systems are possible candidates to be involved in this phenomenon [59]. Actually, previous studies had shown that IDE will interact with 20S proteasome and modulate its proteolytic activity [60]. Our results showed that TC treatment could not only enhance the expression of IDE but also increase the proteasome expressions of 20S (PSMA3) and 19S (PSMC3) subunits. Although the examination was performed for only one subunit of the 20S and 19S particles, respectively, one could not rule out the possibility that TC treatment increased the abundance of proteasome complexes. Further studies are needed to confirm if the proteolytic activity of 20S proteasome is increased by the TC treatment and thus facilitates the degradation of Aβ polypeptides.

It is noteworthy that TC-HT applied after the Aβ administration gives better curative effect than the pretreatment paradigm, which indicates that TC-HT not only has the ability to prevent or protect the neurons against the Aβ-induced cytotoxicity but also could have the potential to cure the disease. The possible explanation for the superior effect of post-treatment could be that some biochemical signals and elevated proteins could interact with the Aβ aggregation simultaneously, and these effects could not maintain until the insult for the pretreated cells. Another advantage is that the post-treatment paradigms for neuronal injury are also more typical in the clinical condition. The present study employed PCR equipment to demonstrate the protective effect of TC-HT in vitro. For in vivo or clinical applications, there are options of other heating devices, such as high-intensity focused ultrasound (HIFU), which has been widely used as a hyperthermal apparatus [61]. The thermal parameters can also be fine-tuned by controlling the heating power and the size of heated volume to meet the specific needs [62].

In summary, this is the first study on the manipulation of stress response by TC-HT, which protects SH-SY5Y human neural cells from H_2_O_2_ and Aβ-induced cytotoxicity. The results confirm that the physical stimulation by TC-HT provides safer and much superior protective effect than continuous HS. The underlying molecular mechanism for the protective effect is partially due to the activation of Akt pathway and the function of some downstream proteins, such as Nrf2, CREB, and HO-1. Further studies are necessary to confirm the involvement of other stress-induced proteins activated by TC-HT in the neuroprotective effect or their ability in degrading or disaggregating the aggregated proteins. It is also advisable for coupling TC-HT with NDD drugs or natural compounds, in order to attain better curative effect for NDD patients.

## Acknowledgments

The authors would like to acknowledge the service provided by the Research Core Facilities 3 Laboratory of the Department of Medical Research at National Taiwan University hospital for use of flow cytometry system.

## Author Contributions

**Conceptualization:** Chih-Yu Chao.

**Data Curation:** Wei-Ting Chen, Yu-Yi Kuo, Guan-Bo Lin, Chih-Yu Chao.

**Formal analysis:** Wei-Ting Chen, Yu-Yi Kuo, Guan-Bo-Lin, Chueh-Hsuan Lu, Hao-Ping Hsu, Yi-Kun Sun, Chih-Yu Chao.

**Funding acquisition:** Chih-Yu Chao.

**Investigation:** Wei-Ting Chen, Yu-Yi Kuo, Guan-Bo Lin, Chih-Yu Chao.

**Project Administration:** Chih-Yu Chao.

**Supervision:** Chih-Yu Chao.

**Validation:** Wei-Ting Chen, Yu-Yi Kuo, Guan-Bo Lin.

**Writing – original draft:** Wei-Ting Chen, Yu-Yi Kuo, Chih-Yu Chao.

**Writing – review & editing:** Chih-Yu Chao.

